# Engineering Outstanding Water Retention and Toughness through Gradient Cellular Architecture and Delignification: A Case Study of Palm Fronds

**DOI:** 10.64898/2026.10.13.758466

**Authors:** Qian Liu, Mohamed Hamid Salim, Ziyong Zhou, Yarjan Abdul Samad, Marco Beaumont, Gustav Nyström, Blaise L. Tardy

## Abstract

Delignification can engineer wood into high-performance multi-scaled materials, building up from its elementary fibrils of nanocellulose across its micro-scaled cellular architecture. However, the potential to engineer other, structurally diverse, non-woody biomass remains largely unexplored. Non-wood plants can add additional hierarchical structures at the mm- and cm-scale. Here, we explore such unique hierarchical architecture as found in date palm fronds, where the vasculature promotes the formation of near-mm scaled vasculature bundles. We also show that along the growth direction, the multi-scale structure varies substantially, i.e. considering the frond base, petiole, and rachis. These tissues are systematically investigated to establish correlations between cellular structure, delignification pathways, and the combination of water dynamics to mechanical performance. A range of property spaces are showcased, highlighting excellent water-storage capacity (∼1212%) in the never-dried porous state and high toughness after air-dry densification. Alkaline, acidic-bleaching, and combined bleach-cold alkali treatments are employed to tailor chemical composition and microstructure selectively. The combined route facilitates nanocellulose liberation and redistribution, generating a hierarchical network that improves moisture retention. These findings establish a structure-processing-property relationship for unconventional plant ultra-structure biomass towards multi-scaled materials with unique features from the nano-to the cm-scales.

## Introduction

Engineered nanostructured wood has emerged as a versatile class of sustainable materials owing to its hierarchical cellular architecture, high strength-to-weight ratio, and tunable functionality [1–5]. By selectively removing lignin and other non-cellulosic components while preserving the aligned cellulose nanofibril network, wood can be transformed into porous frameworks with enhanced water-interaction properties [6,7]. These materials have enabled a broad range of water-management technologies, including passive evaporative cooling [8,9], desalination [10–12], solar evaporation [13,14], atmospheric water harvesting [15,16], and emerging biohybrid systems [17], all of which rely on the precise regulation of water transport, storage, retention, and evaporation within hierarchical cellular architectures. Their performance is fundamentally governed by the hierarchical organization of the cellular framework, which determines water-storage capacity and facilitates efficient capillary-driven transport [18,19]. The interplay between pore-size distribution, interconnected transport pathways, and surface chemistry further regulates water retention and release [20,21]. For applications involving repeated hydration-dehydration cycles, maintaining sufficient mechanical robustness and structural integrity is equally important [22].

Beyond conventional wood, other plant species such as those adapted to arid environments represent an underexplored source of cellular architecture for advanced water-management materials. These plants have evolved specialized tissues to efficiently capture, transport, and retain water under prolonged water-limited conditions, suggesting structural design strategies that differ fundamentally from those of woody species [23,24]. Among them, palm trees, and more specifically date palm trees (Phoenix dactylifera) are particularly attractive because of their multi-scaled cellular organization [25–27]. Unlike conventional wood, which is dominated by relatively homogeneous xylem cells, palm tissues contain abundant parenchyma cells together with vascular bundles that vary substantially in diameter along the plants’ growth axis [28]. Date palm fronds exhibit pronounced spatial variations in cellular architecture from the enlarged frond base attached to the trunk, through the petiole, to the rachis [29]. The basal region possesses a loose, low-density architecture that may favor water storage, whereas the central region is denser and more compact, providing greater mechanical support [30,31]. Furthermore, the high fraction and heterogeneous distribution of parenchyma cells in palm tissues create cellular organizations that are fundamentally different from those found in conventional wood, suggesting opportunities to simultaneously regulate water storage, transport, and mechanical performance [32,33].

Herein, we establish a structure-property framework for date-palm fronds and petioles by linking their tissue-specific cellular architecture to water-management and mechanical performance. The large, loosely packed parenchyma cells of the frond provide abundant lumen space for water uptake and storage, whereas the compact cellular structure and densely distributed vascular bundles of the petiole promote load transfer and compression resistance (Figure 1a). Selective alkaline and bleach delignification further enables these complementary properties to be tailored through controlled modification of tissue composition and porosity (Figure 1b). The resulting bleached frond (Bleached-F) achieves a water-storage capacity of ∼1212% in the never-dried porous state, while the air-dried densified petiole-derived material (Alkaline-P) exhibits a compressive toughness of 18 MJ/m^3^ (Figure 1c). These findings demonstrate how the anatomical diversity of plant tissues can be exploited to transform underutilized date-palm biomass into sustainable materials for water-management and structural applications.

**Figure 1.**
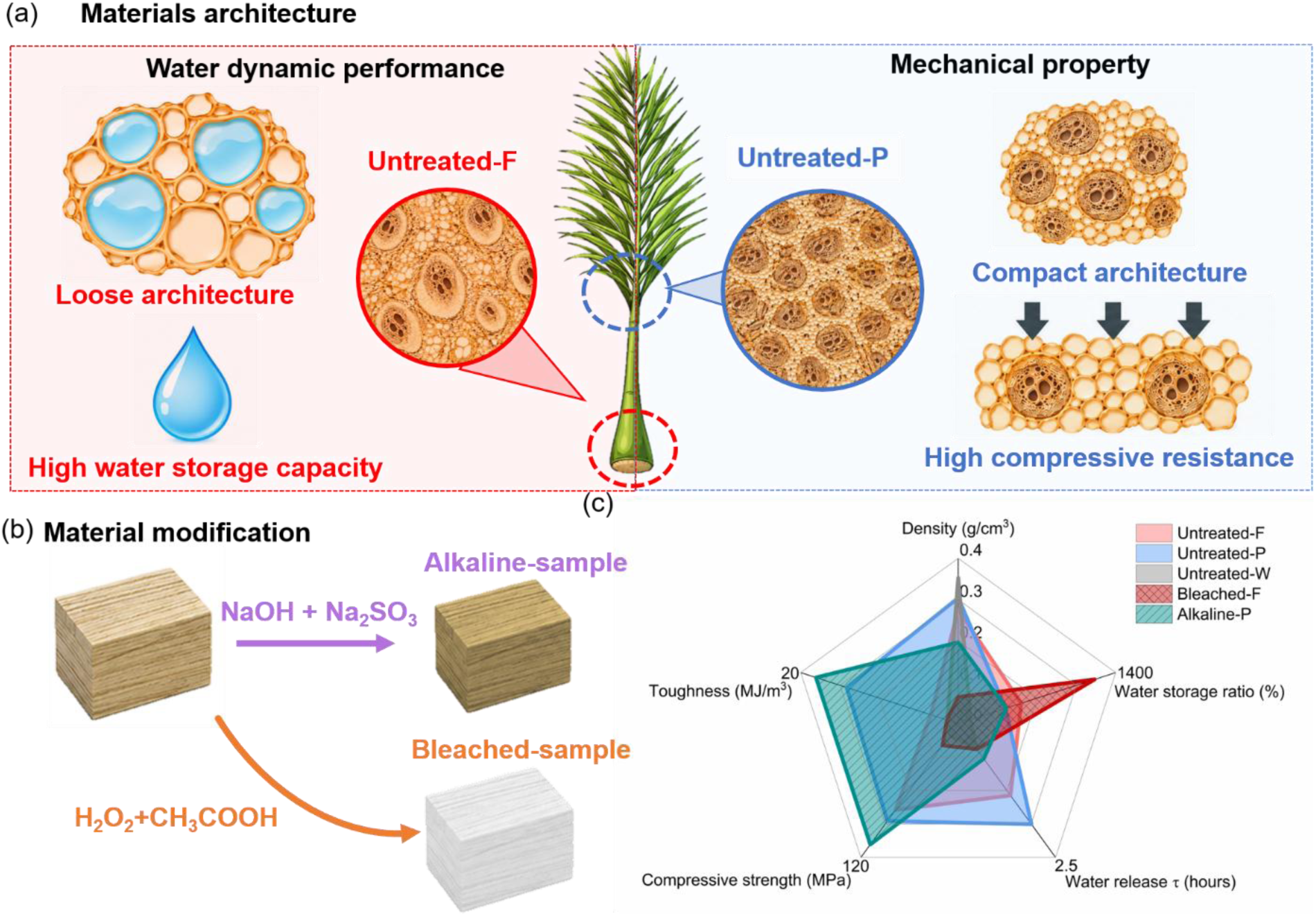
Overview of date palm derived materials. (a) Tissue-specific cellular architectures governing water-storage and compressive performance of untreated frond and untreated petiole. (b) delignification processes for the preparation of delignified samples, and (c) Radar plot summarizing the multi-property performance of untreated and chemically treated samples in comparison with untreated wood. Metrics include density, water storage ratio, water release time constant (*τ*), air-dried maximum compressive strength, and toughness.

## 2. Experimental section

### 2.1 Materials and chemicals

Date palm tree (*Phoenix dactylifera*) samples, including frond (Untreated-F), petiole (Untreated-P), and rachis (Untreated-R) were obtained from local farms in Abu Dhabi - United Arab Emirates and dried at room temperature. Sodium hydroxide (pellets for analysis, CAS No. 1310-73-2), sodium sulfite (powder for analysis, CAS No. 7757-83-7), Sodium chlorite (powder for analysis, CAS No. 7758-19-2), sodium acetate (powder for analysis, CAS No. 127-09-3), hydrogen peroxide (30%, CAS No. 7722-84-1), and acetic acid (anhydrous for analysis, CAS No. 64-19-7) were purchased from Merck KGaA. Deionized water (DI) was used in all the experiments. Balsa wood (natural balsa wood) was purchased from Amazon (Qj-solar) and cut into pieces (designated Untreated-W).

### 2.2 Preparation of delignified date fronds and date petioles

Three untreated date palm tissues (Untreated-F, Untreated-P, and Untreated-R) and a balsa wood reference (Untreated-W) were initially characterized. Only Untreated-F and Untreated-P were selected for subsequent delignification and aerogel preparation. Untreated-W and Untreated-R were evaluated only in their untreated states and were not further processed. The delignification of date palm tree fronds (Untreated-F) and petiole (Untreated-P) was conducted through a range of steps as explained below. For alkaline delignification, Untreated-F and Untreated-P were immersed in a mixed aqueous solution of NaOH (2.5 M) and Na₂SO_3_ (0.4 M) at 90 °C for 6 h. The resulting samples were washed with deionized (DI) water and denoted as Alkaline-F and Alkaline-P, respectively. For acidic-bleaching, Untreated-F and Untreated-P were treated with a peracetic acid solution (PAA) prepared by mixing equal volumes of hydrogen peroxide and glacial acetic acid at 80 °C for 6 h. After thorough washing with DI water, the samples were labelled Bleached-F and Bleached-P, respectively. NaClO2 delignification was selected instead of peracetic acid treatment for aerogel preparation, because it provided a milder lignin removal which was crucial for maintaining the structural integrity (Figure S1a, b). The untreated specimens were first immersed in a NaClO₂ solution buffered to pH ∼4.6 using an acetic acid/sodium acetate buffer and treated at 80 °C for 6h. The delignified samples were then washed with DI water and subsequently soaked in an 8 wt.% NaOH solution at -6 °C for 48 h to obtain aerogel precursors [34]. The samples were repeatedly washed with DI water until a neutral pH was reached to obtain Aerogel-F and Aerogel-P, respectively. All chemical treatments were conducted at a solid-to-liquid mass ratio of 1:25 (w/w). The specimens were cut to approximate dimensions of 1 cm × 1 cm × 1 cm. The sample codes and corresponding treatments are listed in Table 1.

**Table 1:** Treatment Overview of Samples.

| Sample Code | Raw material | Initial Treatment Conditions | Secondary Treatment |
| --- | --- | --- | --- |
| Untreated-F | Date palm frond | - | - |
| Untreated-P | Date palm petiole | - | - |
| Untreated-R | Date palm rachis | - | - |
| Untreated-W | Balsa wood | - | - |
| Aerogel-F | Date palm frond | NaClO<sub>2</sub> (80 °C / 6 h) | NaOH (-6 °C / 48 h) |
| Aerogel-P | Date palm petiole | NaClO<sub>2</sub> (80 °C / 6 h) | NaOH (-6 °C / 48 h) |
| Bleached-F | Date palm frond | H<sub>2</sub>O<sub>2</sub>+CH<sub>3</sub>COOH (80 °C / 6 h) | - |
| Bleached-P | Date palm petiole | H<sub>2</sub>O<sub>2</sub>+CH<sub>3</sub>COOH (80 °C / 6 h) | - |
| Alkaline-F | Date palm frond | NaOH + Na<sub>2</sub>SO<sub>3</sub> (90 °C / 6 h) | - |
| Alkaline-P | Date palm petiole | NaOH + Na<sub>2</sub>SO<sub>3</sub> (90 °C / 6 h) | - |

### 2.3 Characterization

#### 2.3.1 Fourier transform infrared spectroscopy (FTIR)

After drying at 105 °C overnight, the samples were cooled to room temperature and transferred to the FTIR instrument. Spectra were acquired under vacuum using a Bruker Vertex 80v over the range of 4000-400 cm^-1^, with 32 scans and a resolution of 4 cm^-1^. Before the sample measurements, an empty-compartment background spectrum was recorded under the same vacuum conditions and used for background correction. The spectra were subsequently processed using Spectragryph. A third-order Savitzky-Golay smoothing filter was applied using the software-defined interval, and each spectrum was normalized by setting the absorbance of the polysaccharide-associated C-O stretching band at approximately 1060 cm^-1^ to unity.

#### 2.3.2 Thermal properties

The thermogravimetric analysis of the prepared samples was carried out using a thermogravimetric analyzer (TGA Netzsch 449F1) under a dry air atmosphere. Ten different samples (i.e., Untreated-F, Untreated-P, Untreated-R, Untreated-W, Alkaline-F, Alkaline-P, Bleached-F, Bleached-P, Aerogel-F, and Aerogel-P) were heated from 30 °C to 800 °C at a heating rate of 10 °C/min. About 5-10 mg of each sample was used for each analysis.

#### 2.3.3 Microstructure Morphology Analysis

The cross-sectional surface morphologies of Untreated-F, Untreated-P, Untreated-R, Alkaline-F, Alkaline-P, Bleached-F, Bleached-P, Aerogel-F, and Aerogel-P samples were observed via SEM (Quanta 250 FEG) at an accelerating voltage of 5 kV. Two drying protocols were used for specimen preparation. The untreated samples and Aerogel-F/P specimens were freeze-dried before imaging. For Alkaline-F/P, and Bleached-F/P, separate specimens were prepared by either air-drying or freeze-drying to evaluate the influence of the drying method on the observed morphology. All dried specimens were sputter-coated with a thin layer of gold before SEM observation.

Cell areas were calculated using *ImageJ* (National Institutes of Health, USA) software based on the images obtained from SEM, where individual cell regions were measured to quantify cell area for each sample manually. For each sample, cell areas were measured from Figures S2a and 2c for Untreated-F and Untreated-P, respectively. The cell-wall thicknesses of vascular-bundle fibres in untreated frond (Untreated-F) and untreated petiole (Untreated-P) were also manually measured from cross-sectional SEM images using ImageJ. Cell-wall thicknesses were measured from Figures S2b and S2d for Untreated-F and Untreated-P, respectively. The spatial scale of each image was calibrated using the corresponding SEM scale bar. Vascular-bundle fibres were identified as the thick-walled, small-lumen cells surrounding the vascular bundles, whereas vessel elements and larger thin-walled parenchyma cells were excluded. Wall thickness was measured perpendicular to the local wall surface at positions where both wall boundaries were clearly resolved. For each sample, 25 individual wall-thickness measurements were collected, and the results are reported as mean ± standard deviation (n = 25 measurements per sample).

#### 2.3.4 Mechanical Properties

Compression tests were performed using an INSTRON 5948 universal testing machine equipped with a 20 kN load cell in the transversal direction at a crosshead speed of 1 mm/min. This loading rate was selected to maintain a consistently low strain rate despite slight variations in specimen height. The initial contact area of each specimen, corresponding to the loaded surface, was determined by image analysis (*ImageJ* software) and used to calculate the compressive stress. The compressive strain was calculated based on the initial specimen height, which was recorded by the testing machine prior to each test. Before air-drying at room temperature, all specimens had dimensions of 1 cm × 1 cm × 1 cm, and at least five specimens were tested for each experimental condition.

### 2.4 Density

The density of the specimens was determined for both the freeze-dried and air-dried states. The dimensions of each specimen were measured using a digital caliper and *ImageJ* software to calculate its volume, and the mass was determined using an analytical balance. The density was then calculated as the ratio of mass to volume. At least five specimens were measured for each experimental condition.

### 2.5 Water Retention and Dynamics

Specimens of Untreated samples and delignified samples (Alkaline-F/P, Bleached-F/P, and Aerogel-F/P), each with an approximate dimension of 1 cm × 1 cm × 1 cm, were used for the measurements. The untreated and delignified specimens (never dried state) were maintained in DI at least for 7 days before testing to make sure all samples are in water-saturated state, and the water-saturated mass (*m*_1_) was recorded.

For the water-retention test, the saturated specimens were dried at 50 °C for 8 h, and their mass, *m*_2_(*t*), was recorded at 1 h intervals. After the retention test, the specimens were further oven-dried at 105 °C to a constant mass to determine their dry mass (*m*_3_). At least five independent specimens were measured for each sample condition. The water-storage ratio was calculated as:

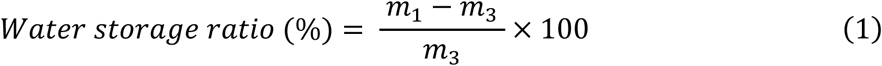

Water-retention performance was quantified using the water-release time constant (*τ*) [35], operationally defined as the time required for 63.2% of the initially stored water to evaporate. The specimen mass corresponding to this point, (*m*^∗^), was calculated as:

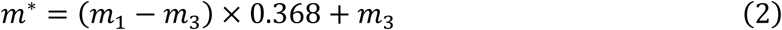

Because the specimen mass was recorded at discrete time intervals, (*τ*) was determined by linear interpolation:

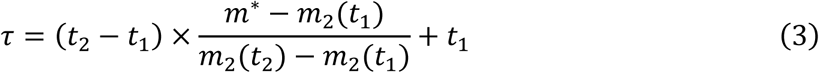

where *t*_1_ and *t*_2_ are the consecutive measurement times immediately before and after the specimen mass crossed *m*^∗^, respectively. The corresponding measured masses are *m*_2_(*t*_1_) and *m*_2_(*t*_2_)

## 3. Results and discussion

### 3.1 Architecture and mechanical properties of untreated date palm samples and wood sample

The untreated frond, petiole, rachis, and wood samples, hereafter denoted as Untreated-F, Untreated-P, Untreated-R, and Untreated-W, respectively, were analysed to evaluate their native cellular architectures and physical properties. Cellular morphology analysis revealed that Untreated-F and Untreated-P possessed distinct tissue architectures characterized by a heterogeneous distribution of vascular bundles embedded within highly porous parenchymatous matrices (Figure 2a). Compared with Untreated-F, where the vascular bundle (2a1) had diameters reaching mm-scale (Figure S2a), the diameter of Untreated-P vascular bundles typically no more than 0.5 mm (Figure 2b, c). This indicates a narrowing of cellular structures from the base of the frond to its end, which may result in a denser microstructure and thus enhanced mechanical properties. In addition, the average vascular cell wall thicknesses of Untreated-F (Figure S2b) and Untreated-P (Figure S2d) were 1.0±0.4 μm and 0.85 ± 0.2 μm, respectively. These values were lower than the typical cell wall thicknesses reported for wood fibres (∼1.5 μm) [36] and vessel elements, which generally vary depending on species, tissue type, and growth conditions [37]. Other features observed along the vessel wall included micro-cracks and large openings in Untreated-P, which originated from natural pit apertures that facilitate fluid exchange between adjacent cells (Figure S2d).

**Figure 2.**
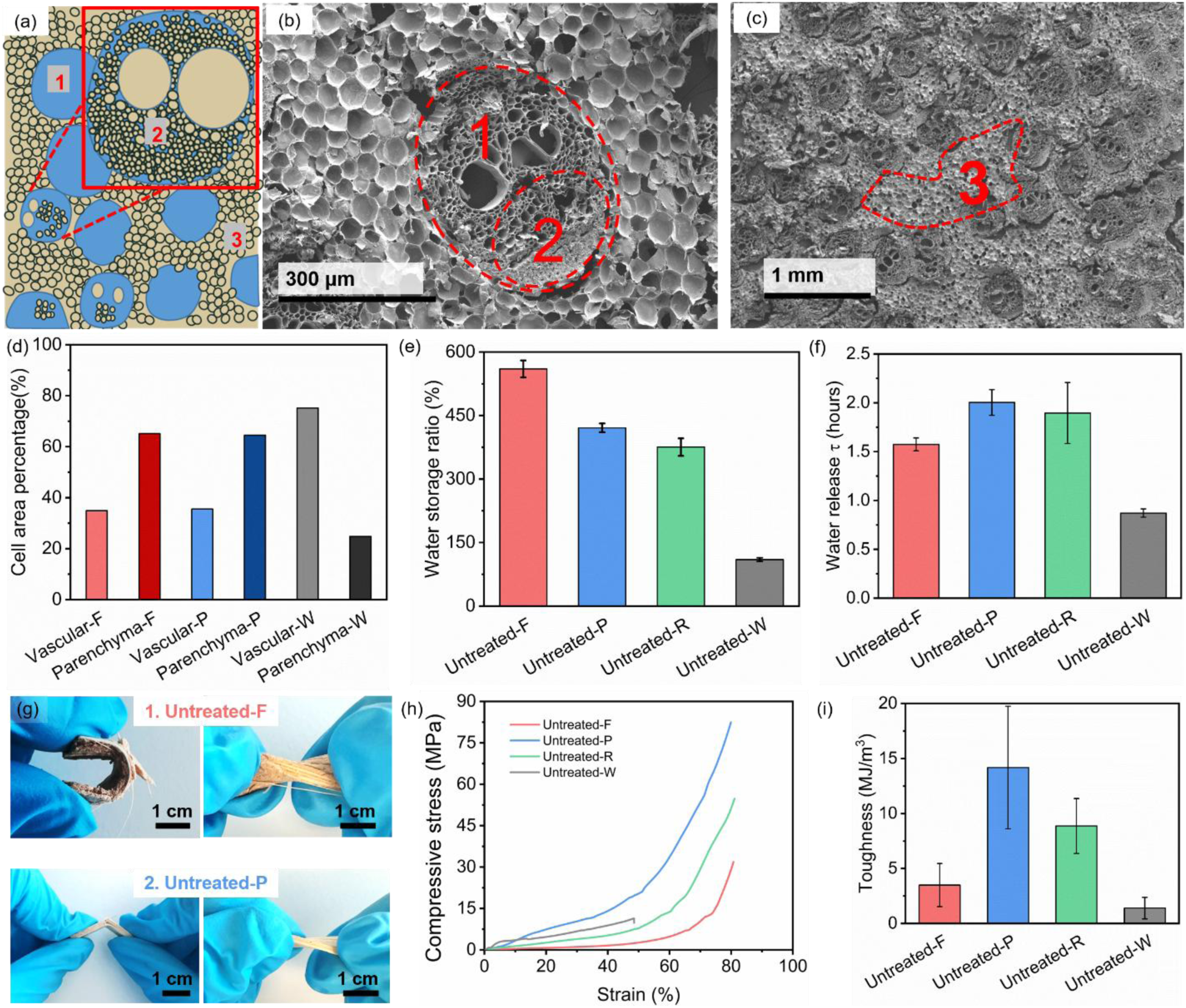
(a) Schematic illustration of the major cells in date palm-derived samples. SEM images of Untreated-P at different magnifications are shown in (b, c). (d) Quantitative analysis of the area fraction of the major cells in Untreated-F, Untreated-P, and Untreated-W. (e) Water storage ratio and (f) water release time constant of the untreated samples. Mechanical deformation behavior under manual bending of (g) Untreated-F and Untreated-P. (h) Representative transversal compressive stress curve and (i) average compressive toughness of the untreated date palm-derived and wood samples.

The cell area ratios of vascular bundle to parenchyma cells in Untreated-F (Figure S2a), Untreated-P (Figure 2c), and reported balsa wood [38] were quantitatively analyzed (Figure 2d). The ratio of parenchyma cells to the total cell area in both Untreated-F (65%) and Untreated-P (64%) is substantially higher than that in natural wood (24.8 %). The higher proportion of parenchyma cells likely contributes to the enhanced water transport, storage, and retention capacity of the date palm-derived samples. This may also be associated with their faster growth rate, compared with wood.

The impact of cellular architecture on water storage as well as release dynamics was evaluated. Untreated-F with a lightweight framework exhibited the maximum water storage ratio reaching 560% (5.6-fold its own dried weight), which is 5.1 times higher than that of Untreated-W (herein high-density balsa wood, Figure 2e). Untreated-P and Untreated-R exhibited similar water storage ability with a storage ratio of 420% and 375%, respectively, corresponding with their more compact cell architecture compared with fronds (Figure S3a). The time required for 63.2% of the stored water to evaporate was evaluated as a release constant for each of the materials (Figure 2f), following the conventional definition of the time constant for a first-order exponential. The best water retention performance was observed in the Untreated-P, at a time (τ) of 2 h, which was double that of Untreated-W, highlighting possible combination of high water storage and retention associated with the cellular architecture of petioles, where both frond and rachis samples also outperformed wood.

The impact of cellular architecture on mechanical properties was evaluated. Surprisingly, a higher mechanical deformability of date palm-derived materials under bending and torsional loading was observed than untreated wood (Figure 2g, S4). Under bending (under a radius of curvature of ∼1.5 cm), Untreated-F (Figure 2g top-left, Figure S4a) maintained structural integrity, indicating excellent flexibility and resistance to brittle failure, which was not observed in the other untreated samples (Figure S4b-d). When subjected to torsional deformation, Untreated-F (Figure 2g, top-right) and Untreated-P (Figure 2g, bottom-right) sustained significant twisting (∼45⁰ cm^-1^) without fracture. These results suggested that variations in cellular architecture, particularly cell size distribution and structural heterogeneity, may play a critical role in accommodating multidirectional stress and thus overall mechanical resilience. Accordingly, the compressive stress performance of air-dried samples in transversal direction was tested to further explore their microstructure impact on energy absorption. All samples showed a characteristic compressive response consisting of an initial elastic region, followed by progressive cell-wall collapse and a densification stage at high strains (Figure 2h), typical of cellular materials. These regimes are dependent on cell diameter, cell wall thickness, and the relative size distribution of these cells [39]. Untreated-P exhibited a maximum load of 5040 N and a compressive stress of about 80 MPa at 80% strain, followed by Untreated-R (∼55 MPa), and Untreated-F (∼32 MPa) at the same strain. The lowest compressive strength was observed in Untreated-W (∼11 MPa), which fractured early at a strain of around 45%. Untreated-P carried load effectively at relatively low strains and entered the stiffness-increase and densification stages earlier, suggesting a cellular structure more favorable for load transfer. Untreated-F initially underwent cell-wall bending, progressive pore collapse, followed by densification at approximately 75% strain, consistent with a more porous structure in Untreated-F.

The corresponding toughness values (Figure 2i) showed a similar trend, demonstrating significant differences in energy absorption capacity among the samples. Untreated-P exhibited the highest average toughness (∼14.2 MJ/m^3^), which was approximately four times greater than that of Untreated-F (∼3.5 MJ/m^3^), and more than an order of magnitude higher than that of Untreated-W (∼1.4 MJ/m^3^). These results indicate that Untreated-P possesses the greatest ability to sustain deformation while absorbing mechanical energy, whereas Untreated-W is considerably more brittle and susceptible to compressive failure. The superior compressive performance of the Untreated-P may be attributed to its higher density (Figure S3b) in air dry state and more importantly to its cellular architecture (Figure 2d).

In summary, the date palm derived samples exhibit several distinctive structural and functional characteristics: (i) a cellular architecture consisting of parenchyma cells and vascular bundles, with the parenchyma cells accounting for approximately 65% of the cross-sectional area, nearly three times the fraction observed in wood; (ii) this unique cellular organization results in a more open microstructure, thereby enhancing water storage and retention capacity; and (iii) these structural characteristics contribute to superior energy absorption and mechanical resilience, yielding exceptional compressive toughness, particularly in the Untreated-P fraction of the palm-tree tissues, which is nearly ten times greater than that of untreated wood.

### 3.2 Chemical and physical properties of delignified date palm samples

Delignification, and more holistically the removal of non-cellulosics, plays an important role in the end properties of plant materials [40]. To evaluate the potential of non-cellulosics removal to improve the inherent properties of date frond and petioles, three different routes were used to prepare delignified frond and petiole (Figure 3a): (i) alkaline delignification using the combination of NaOH and Na_2_SO_3_ to remove lignin, also resulting in additional chromophores [41] that made the samples appear darker (Figure 3a1 - *Alkaline*); (ii) Acid bleach delignification using peracetic acid which resulted in a white material (Figure 3a2-*bleached*); and (iii) the combination of bleach (NaClO2) and cold alkali dissolution process achieving both lignin removal and cellulose migration from the cell wall of date frond and petiole into the cell’s lumen (Figure 3a3 - *Aerogel*). Of note, as the untreated rachis did not outperform either the petiole or the frond base in the targeted properties, it was not evaluated for delignification and subsequent properties evaluation.

**Figure 3.**
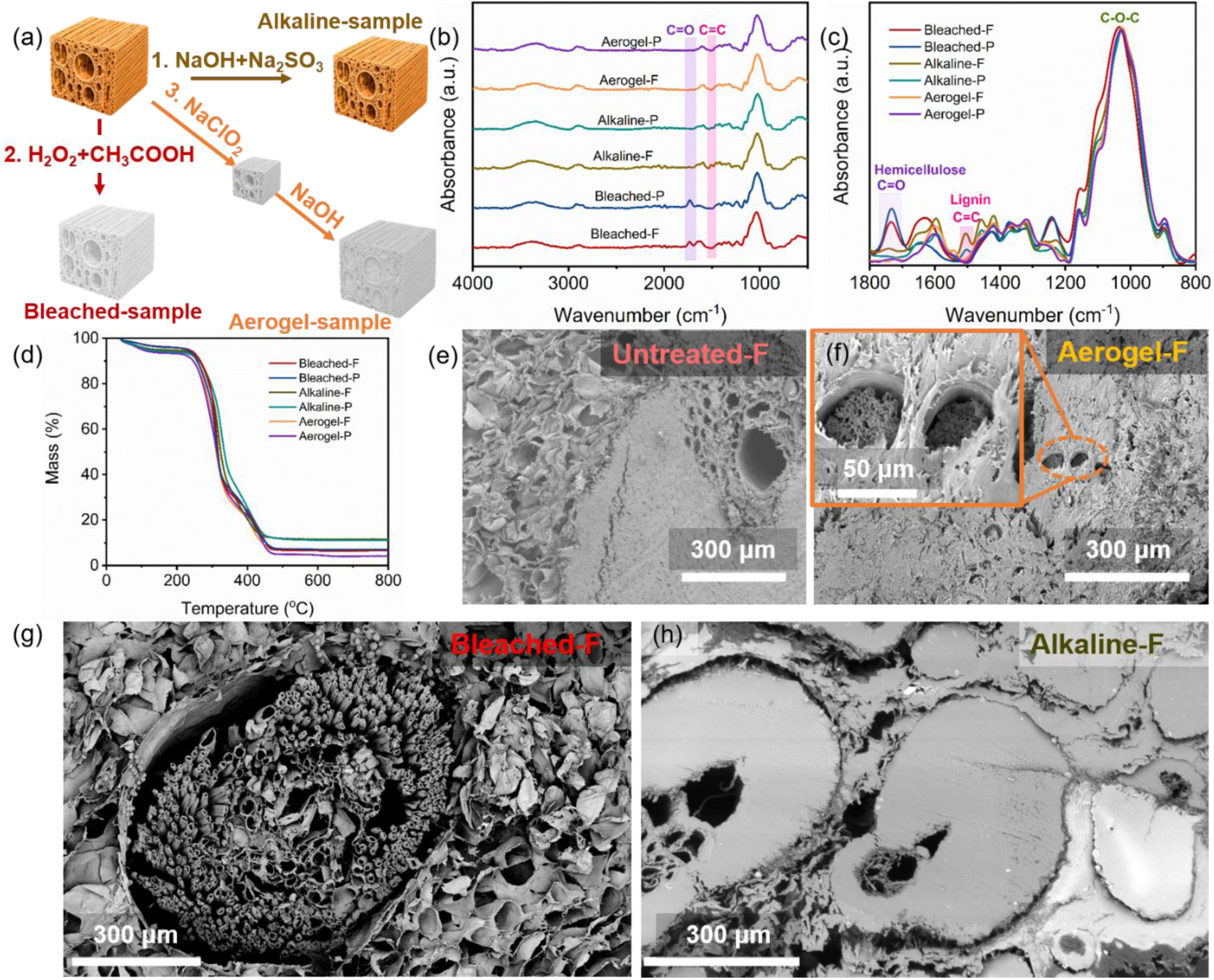
Preparation routes, chemical characterization, and microstructure of delignified samples. (a) Schematic illustration of the three delignification routes used to prepare delignified frond and petiole samples. (b, c) Fourier transform infrared (FTIR) spectra and (d) thermogravimetric analysis (TGA) of the delignified samples. SEM images of (e) Untreated-F, (f) freeze-dried frond aerogel, (g) Bleach-treated frond after freeze drying, and (h) alkaline-treated frond after air drying.

The loss of the band near 1740 cm^-1^ from FTIR (Figure 3b, c) is consistent with changes in hemicellulose-associated ester groups, including deacetylation and possible hemicellulose removal by Alkaline treatment [42]. Compared with the alkaline-treated samples, both bleach-treated frond and petiole exhibited a relatively lower intensity of the lignin-associated band at approximately 1510 cm^-1^ (Figure 3c). This decrease was consistent with more extensive removal or bleach modification of aromatic lignin structures during bleach delignification [43,44]. Bleached-P retained a stronger band at approximately 1740 cm^-1^ than Bleached-F, which may suggest a greater abundance of residual hemicellulose-related structures, consistent with the compositional differences observed between the corresponding untreated tissues (Figure S1d, Supplementary Discussion 1).

Thermogravimetric analysis (TGA) of the treated date palm-derived samples revealed multi-stage degradation behavior with noticeable differences induced by chemical treatments. The untreated and treated date-palm tissues exhibited distinct thermogravimetric profiles. All samples showed a small initial mass loss below ∼150 °C, mainly associated with the removal of physically adsorbed water, followed by the principal degradation of the lignocellulosic components at around 220-430 °C [45]. Among the untreated samples (Figure S1e, Supplementary Discussion 2), date palm derived samples displayed relatively broad and comparable degradation profiles, whereas Untreated-W maintained its mass to a somewhat higher temperature before undergoing a more abrupt mass loss. Nevertheless, Untreated-W exhibited the lowest residual mass at 800 °C (∼15%), compared with ∼25-30% for the date palm derived samples.

After delignification processing, the degradation profiles of the frond- and petiole-derived samples became substantially more similar. This convergence suggests that the delignification reduced the compositional differences between the Untreated samples and may produce more compositionally uniform, cellulose-rich scaffolds [46]. Delignified samples also showed markedly lower residual masses at 800 °C (4-11%) than the untreated samples, consistent with the removal of lignin and other components that contribute to high-temperature residue formation. In addition, compared to bleach-treated samples, the relatively high residual mass was observed for alkaline-treated samples. The aerogel samples exhibited the lowest final residual masses, which may reflect treatment-induced changes in the lignocellulosic composition and a reduced inorganic fraction.

The microstructures of delignified samples were observed by scanning electron microscopy (SEM) of freeze-dried and air-dried samples to investigate the impact of the chemical treatment as well as drying processes. For freeze-dried samples, Aerogel-F (Figure 3f) and Aerogel-P (Figure S5b) exhibited fibrillar structures both on the exposed macrofibrillar surfaces and within the cell lumina where they formed loose yet continuous networks, which were not observed in Untreated-F. Bleach treatment (Figure 3g) produced a relatively open cellular architecture, with Bleached-F and Bleached-P exhibiting the lowest apparent densities (Figure S6a), which could be beneficial for water storage. In contrast, alkaline treatment (Figure S5c) induced more pronounced cell wall degradation and fibrillation, resulting in disconnected pores. Under each treatment, the petiole-derived sample exhibited a higher apparent density than its frond-derived counterpart, consistent with the more compact native architecture of the petiole.

In contrast to freeze-dried samples, the air-dried samples were densified and exhibited severe structural collapse and densification for Bleached-F (Figure S5d) and Alkaline-F (Figure 3h). The originally open cellular network was no longer distinguishable with most lumens compressed or completely closed for Bleached-F and Alkaline-F. The cell walls appeared flattened and fused together, producing a compact and relatively smooth surface. Similar microstructural evolution was observed in the petiole-derived samples. Alkaline-P (Figures S5e) and Bleached-P (Figures S5f) show compact and partially collapsed cellular structure and they retain a recognizable cellular morphology. The cell walls are relatively thick and continuous, forming a dense load-bearing framework with limited intercellular voids. In addition, a denser microstructure of petiole-derived samples (Figures S5f) is observed than the corresponding frond-derived sample (Figure S5d) due to their distinct microstructure even when treated by the same processes.

### 3.3 Mechanical performance, water storage, and water dynamics of delignified date palm samples

Delignification generates a highly porous structure and thus low density (Figure S6a) of samples treated by freeze drying, which can enhance water storage and retention ability of delignified samples. Compared with delignified petiole samples, frond samples subjected to the same delignification treatment exhibited higher water storage ratio (Figure 4a). Bleached-F showed the highest water storage ratio (1212%) among all samples, which may be associated with its low density (∼0.06 g/cm^3^). In contrast, alkaline-treated specimens showed the lowest water storage ratio for both delignified frond and petiole samples.

**Figure 4.**
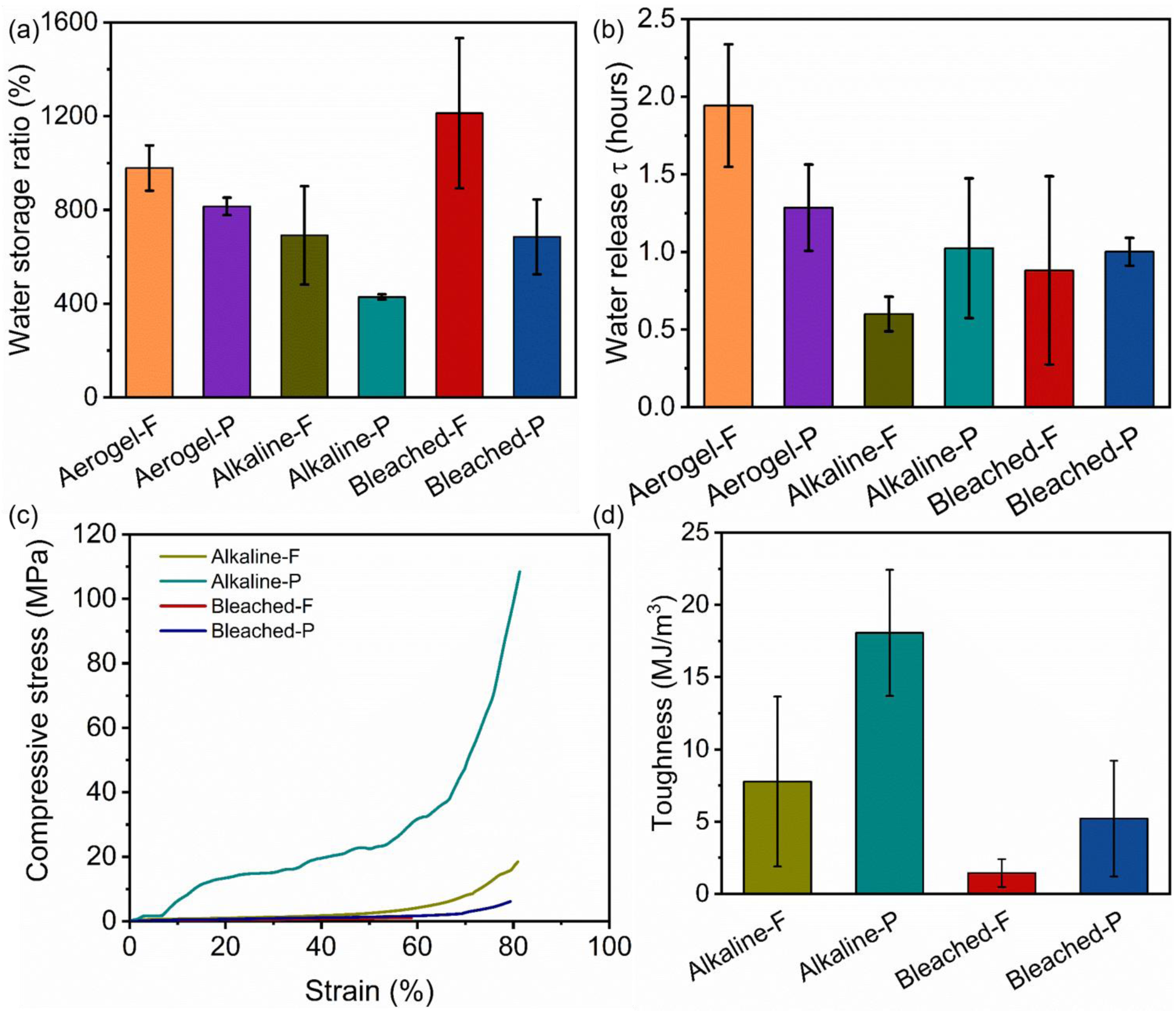
Water management behavior and compressive mechanical performance of delignified samples. (a) Water storage ratio, (b) water release time constant (*τ*), (c) representative compressive stress-strain behavior, and (d) compressive toughness.

The water release time constant (Figure 4b) post-treatment was then evaluated across samples. Aerogel samples (Aerogel-F and Aerogel-P) exhibited superior water retention compared with other delignification-treated samples, likely due to the re-deposited cellulose fibers within the cell lumen that hinder water release (Figure 3f, S5b). Compared with their corresponding untreated samples, Aerogel-F showed a time constant of 1.94 h which was even higher than that of Untreated-F (1.57 h). In contrast, the other treatments resulted in reduced water retention performance. A similar comparison was also made with the petiole-derived samples. The reduced water-retention capacity after delignification treatment may be associated with the partial collapse of the micro- and nanostructures (Figure 3g, S5c). Additionally, the time constants of Alkaline-P and Bleached-P were slightly higher than those of Alkaline-F and Bleached-F, respectively, which may be attributed to their denser structure evidenced by higher freeze-dried density (Figure S6a).

To investigate the impact of delignification on the mechanical properties of date frond and date petiole, the compressive toughness and strength of the air-dried delignified samples were measured across the transversal direction. Among the treated samples, Alkaline-P exhibited the highest compressive strength (∼108 MPa at 81.3% strain). Following initial cell-wall bending, its dense vascular-bundle network enabled efficient load transfer and progressive pore collapse, with densification beginning at around 50%. In comparison, Alkaline-F underwent an extended cell-wall bending and pore-collapse stage, followed by densification at ∼70%, reaching around 18 MPa at 80.9% strain (Figure 4c). Bleach treated samples displayed a prolonged low-stress collapse and weak densification, indicating that bleach treatment substantially weakened their cell walls and intercellular load-bearing network. These differences are closely associated with samples’ air-dried microstructure (Figure S5 d-f) and density (Figure S6b). Alkaline-P exhibited the highest average air-dried density (∼1.0 g/cm^3^) as well as compact microstructure (Figure S5e), whereas Bleached-F had the lowest density (∼0.2 g/cm^3^) and showed the poorest mechanical performance. The higher density reflects a greater volume fraction of load-bearing cell-wall material and a more compact cellular architecture, thereby enhancing the material’s resistance to compressive deformation.

The toughness measurements exhibited a similar trend, confirming significant differences in energy absorption capacity after delignification. Alkaline-P displayed the highest compressive toughness (approximately 18 MJ/m^3^) and Bleached-F exhibited the lowest toughness. Compared with bleach treatment, the alkaline treatment enabled the samples to sustain larger compressive deformation while maintaining a higher load-bearing capacity, resulting in substantially greater energy absorption. The positive correlation between air-dried density and compressive performance further suggests that preserving the cellular framework during delignification is critical for maintaining mechanical integrity.

These findings highlight that (i) delignification treatments produced distinct property profiles with bleaching increased water-storage capacity, whereas alkaline treatment yielded higher compressive energy absorption in air-dried specimens. Most treated samples released water faster than their untreated counterparts while Aerogel-F was the exception; (ii) petiole-based samples consistently exhibit higher mechanical strength than frond-based counterparts under identical delignification conditions, owing to their distinctive intrinsic microstructure.

### 3.4 Comprehensive comparisons of prepared materials’ properties

Ashby maps were employed to comprehensively compare the unique properties of the prepared delignified date frond and date petiole samples with untreated materials, focusing on water dynamic behavior, and mechanical performance. The untreated samples included Untreated-F, Untreated-P, Untreated-R and Untreated-W, while the prepared samples comprised Alkaline-F/P and Bleached-F/P as well as Aerogel-F/P.

A clear inverse structure-function relationship between freeze-dried density and water storage ratio in the prepared materials is revealed in Figure 5a, demonstrating that microstructural porosity governs water storage performance. Samples with low density (∼0.07-0.15 g/cm^3^), corresponding to more open and highly porous cellular architectures, exhibit markedly enhanced water storage capacities (∼700-1212%), while denser structures (>0.25 g/cm^3^) show significantly reduced water storage ratio (<500%). This trend indicates that delignification can increase accessible pore volume and capillary pathways, thereby promoting water absorption. In addition, the spread in water storage at similar densities suggests that beyond bulk density, cell types and distribution also contribute to water dynamics. For instance, the water storage ratio of Untreated-F (∼560%) is higher than that of Alkaline-P (428%), even though the density of Untreated-F is greater than Alkaline-P. Therefore, the delignification process and the intrinsic microstructure of the material govern its hydration behavior, underscoring how the precursor architecture and processing conditions collectively tailor water-related performance.

**Figure 5.**
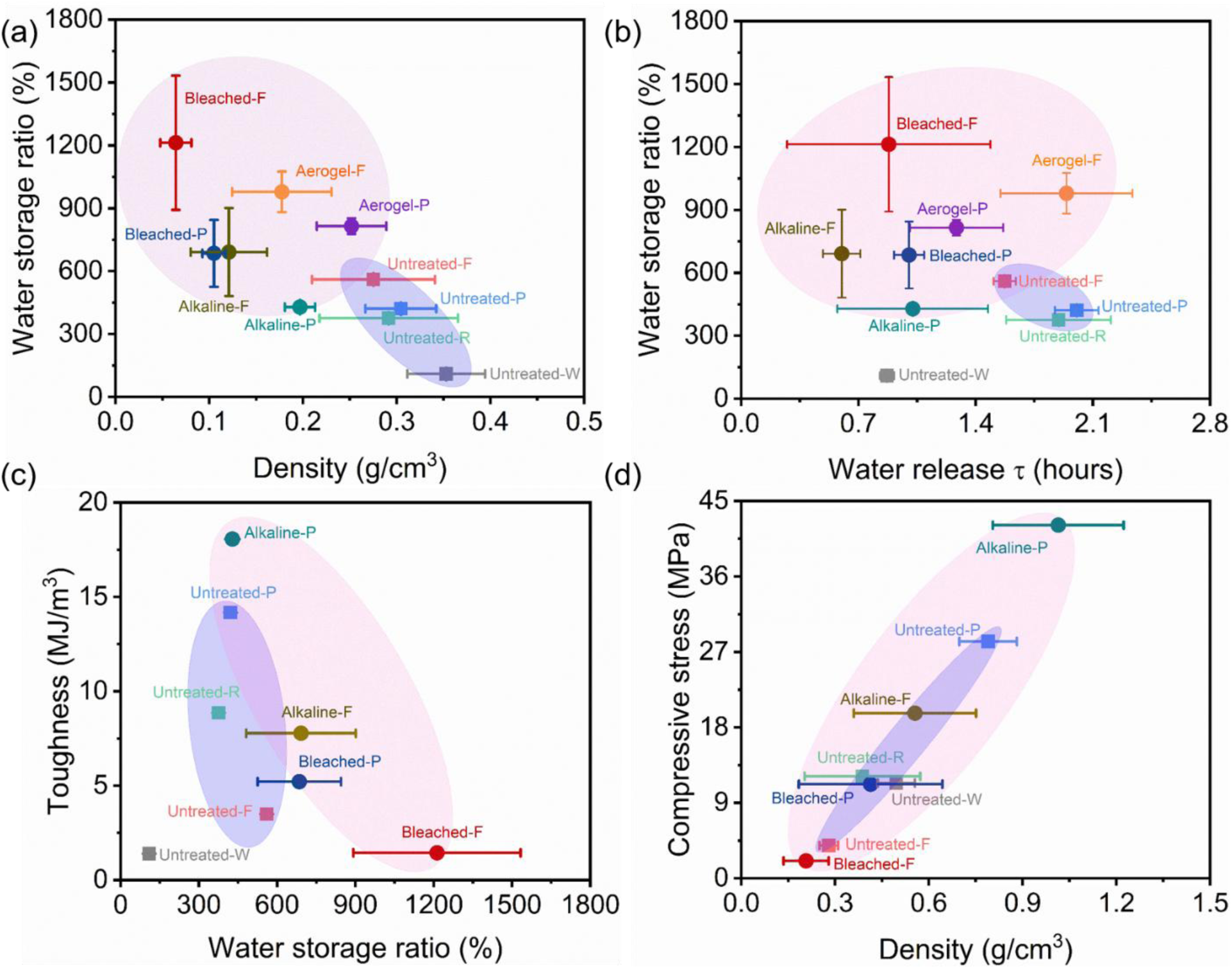
Ashby maps comparing the performance of delignified date palm samples (date frond and date petiole) with untreated materials. Water storage ratio versus (a) freeze-dried density, and (b) water release time constant (*τ*), (c) mean air-dried toughness versus water storage ratio, and (d) mean compressive strength versus air-dried density. All samples are distinguished by a consistent color scheme across panels.

Water dynamic behavior (Figure 5b) further distinguishes the microstructure impact on prepared date palm derived samples and untreated wood. Untreated date palm-derived samples generally demonstrate superior water storage performance with better water retention ability compared with Untreated-W. Chemical treatments affected the water-retention behavior of the frond-derived and petiole-derived samples. Samples with improved water storage ratio by delignification treatment typically exhibited lower water-release time constants than their corresponding untreated samples, indicating faster water release, e.g., Aerogel-P, Bleached-P, and Alkaline-P exhibited water release time constant decrease of 36%, 50%, and 49% compared with Untreated-P, respectively. Aerogel-F was the only exception, showing a longer water-release time than Untreated-F. This behavior may be associated with the loose fibrillar network observed within the cell lumina of Aerogel-F (Figure 3f), which could enhance capillary interactions and delay water release.

The relationship between water storage ratio and air-dried toughness (Figure 5c) of samples reveals an overall trade-off between hydration performance and mechanical strength. In general, samples with high water storage ratio tend to exhibit low toughness after air-dry densification, indicating that the structural modifications promoting water uptake compromise the material’s ability to dissipate mechanical energy. However, this relationship is strongly influenced by the delignification strategy. Alkaline-delignified samples deviate positively from the overall trend, maintaining relatively high toughness despite increased water storage capacity. Alkaline-P exhibited the highest mean toughness and maintained a water storage ratio slightly higher than the Untreated-P, suggesting alkaline treatment appears to preserve water storage capacity without sacrificing toughness, thereby mitigating the apparent trade-off observed for the other treatment routes. In contrast, increased water storage following acidic bleaching was accompanied by a pronounced loss of toughness. This behavior implies that bleach treatment induces more severe degradation of the cell wall framework, weakening the structural integrity required for effective energy dissipation. Overall, the results demonstrate that water storage capacity and toughness are generally competing properties, but their balance is governed by the delignification mechanism, highlighting the critical role of precursor microstructure in simultaneously optimizing hydration and mechanical performance.

The relationship between air-dried density and average compressive stress (Figure 5d) demonstrates a positive correlation, indicating that densification is the primary factor governing the load-bearing capacity of the compressed materials. To enable a consistent comparison among samples with different failure behaviors, the average compressive stress at 60% strain was used for all samples except Untreated-W that fractured at approximately 45% strain (Figure S3c). Although the untreated samples remained intact beyond 80% strain, the delignified samples generally failed at around 70% strain. Therefore, stress at 60% strain provides a representative metric for comparing date palm derived samples’ compressive performance, while for Untreated-W the strain was selected at 45%. High-density samples exhibit great compressive strength, with Alkaline-P achieving both the highest density and the highest compressive strength. In addition, the delignification strategy also significantly influences this relationship. Alkaline-treated samples exhibit higher compressive strength than their untreated counterparts at comparable densities, suggesting that alkaline delignification preserves a more efficient load-bearing cellulose framework during delignification. In contrast, bleach delignification markedly reduces the compressive strength, particularly for Bleached-F. This pronounced decrease in compressive stress of bleach-treated samples indicates that bleach treatment compromises the structural integrity of the cell wall network, thereby limiting its ability to sustain compressive loads.

Overall, the Ashby analysis demonstrates a correlation between density, compressive strength, and water storage ratio. While water retention remains uncorrelated to water storage ratio, highlighting distinct mechanisms associated with each plant tissue and pretreatment chemistry. The prepared palm-derived materials occupy a unique materials space distinct from conventional wood systems. They combine low density with enhanced water storage ability and retention capabilities while maintaining superior load-bearing performance compared with natural wood. Delignification further enhances both the water management properties and the mechanical performance of the date palm-derived materials.

## Conclusion

In this work, we systematically elucidate the structure-property relationships associated with the impact of delignification for a plant-based material with ultrastructures extending beyond that of wood. By doing so, we highlight that tissue structures with heterogeneous cellular architectures can lead to new property spaces. These tissues possess a distinctive cellular architecture comprising vascular bundles and parenchyma cells, with a parenchyma area fraction of approximately 65%. In the freeze-dried state, this cellular architecture provides a porous framework associated with enhanced water storage and relative water retention compared with freeze-dried untreated wood. Separately, after air-dry densification, the date palm-derived materials exhibit greater compressive strength and toughness than the corresponding air-dried untreated wood. Delignification further tailors both the chemical composition and microstructure of these date palm derived materials. Bleach delignification produces highly porous architectures that favor exceptional water storage capacity (∼1212%), whereas alkaline treatment better preserves the cellulose framework, resulting in high compressive strength after air-drying. Under identical delignification conditions, petiole-based materials consistently outperform frond-based counterparts in mechanical strength, underscoring the critical role of native microstructural differences. Overall, this study highlights the versatility of delignification-induced structural engineering of non-wood plant tissues and establishes palm biomass and other related plant ultrastructures as a promising and sustainable platform for advanced functional materials, with potential applications in absorbents, water management systems, and high mechanical strength components.

## Acknowledgements

We acknowledge funding support by the Khalifa University of Science and Technology (KUST) Faculty Startup Project (Project code: 84741140-FSU-2022-021), and the Food Security and Technology Center (FSTC), through KU-FSTC, Project ID: KU-INT-FSTC-2026-8474000576. The authors gratefully acknowledge Abdulla bin Tamim for providing the raw materials used in this research.

