## Supplemental Files for "Engineering Outstanding Water Retention and Toughness through Gradient Cellular Architecture and Delignification: A Case Study of Palm Fronds"

1. Supplementary Figure

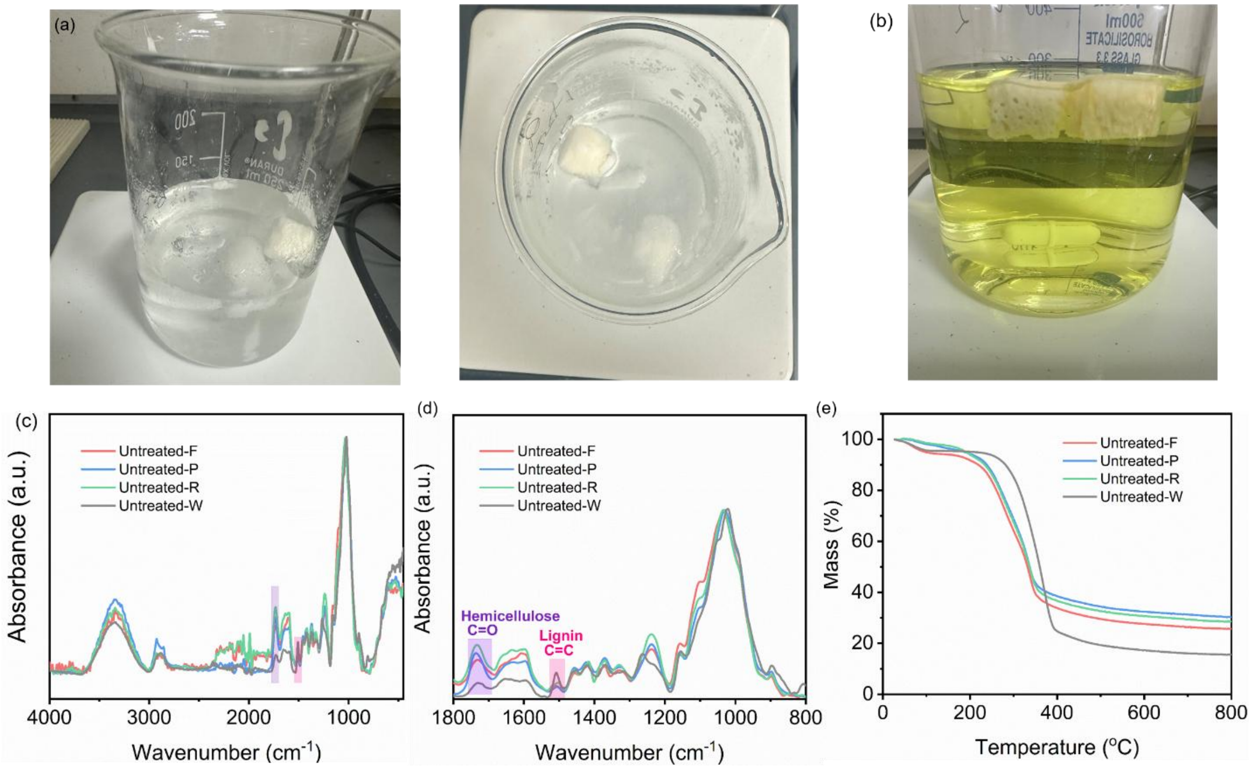

Figure S 1. Photographs of bleach delignification of date palm frond using (a) peracetic acid and (b) $NaClO_2$ after 2 h of treatment. (c, d) Fourier-transform infrared (FTIR) spectra and (e) thermogravimetric analysis (TGA) properties of untreated date palm-derived and wood samples.

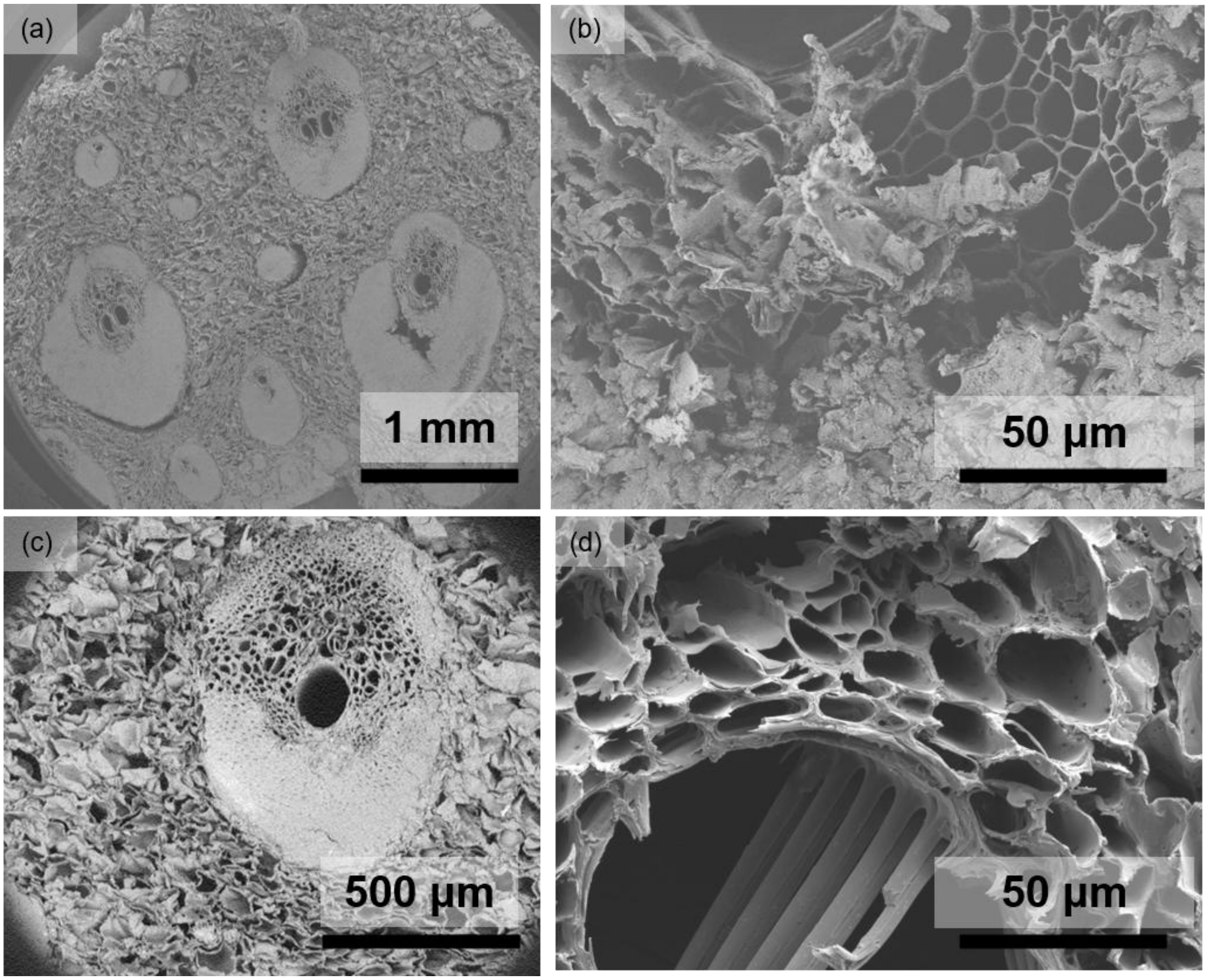

Figure S 2. SEM of (a, b) Untreated-F, and (c, d) Untreated-P.

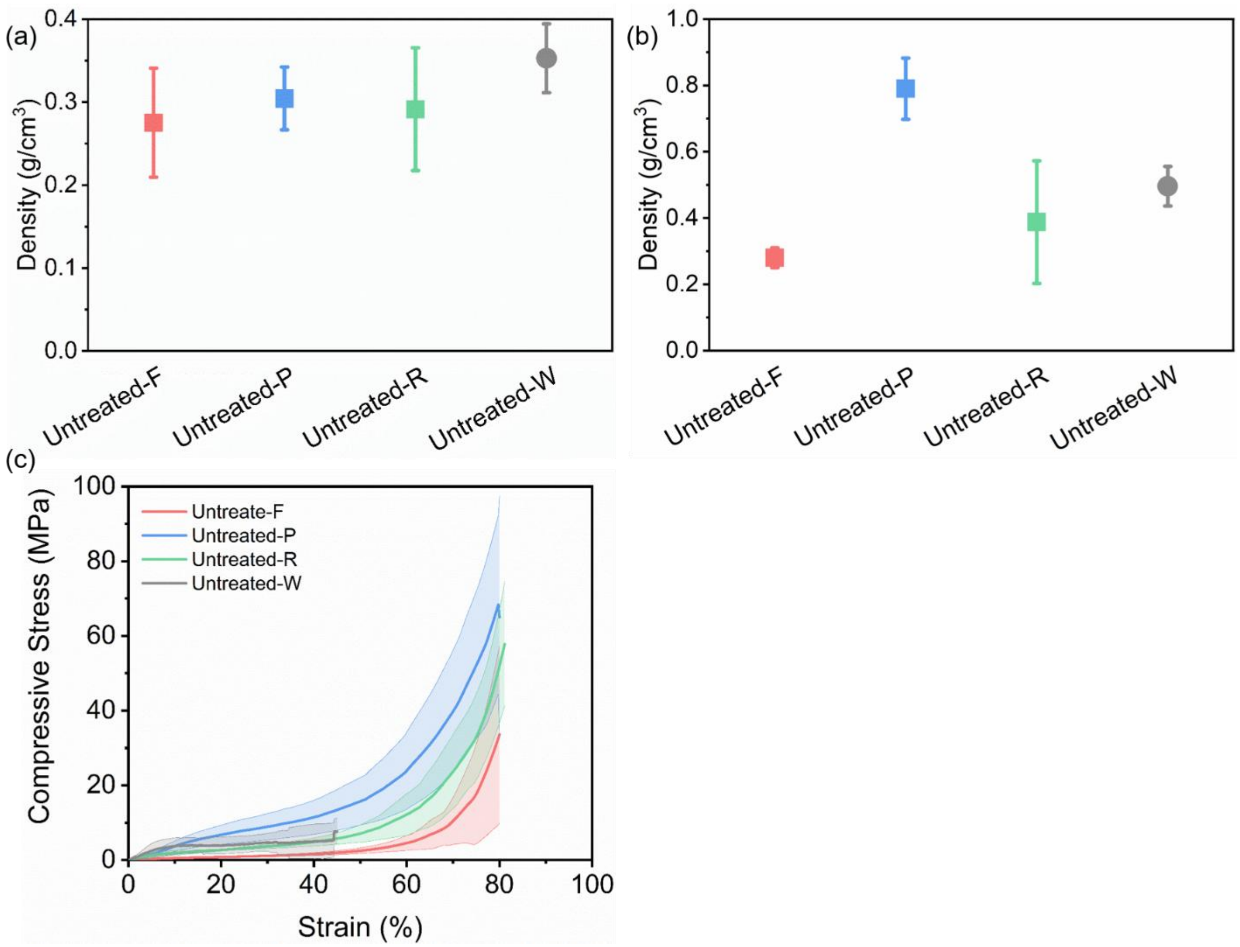

Figure S 3. Density of Untreated samples by (a) freeze-drying, (b) air-drying, and (c) transversal compressive strength representative curve of untreated samples under air-drying condition.

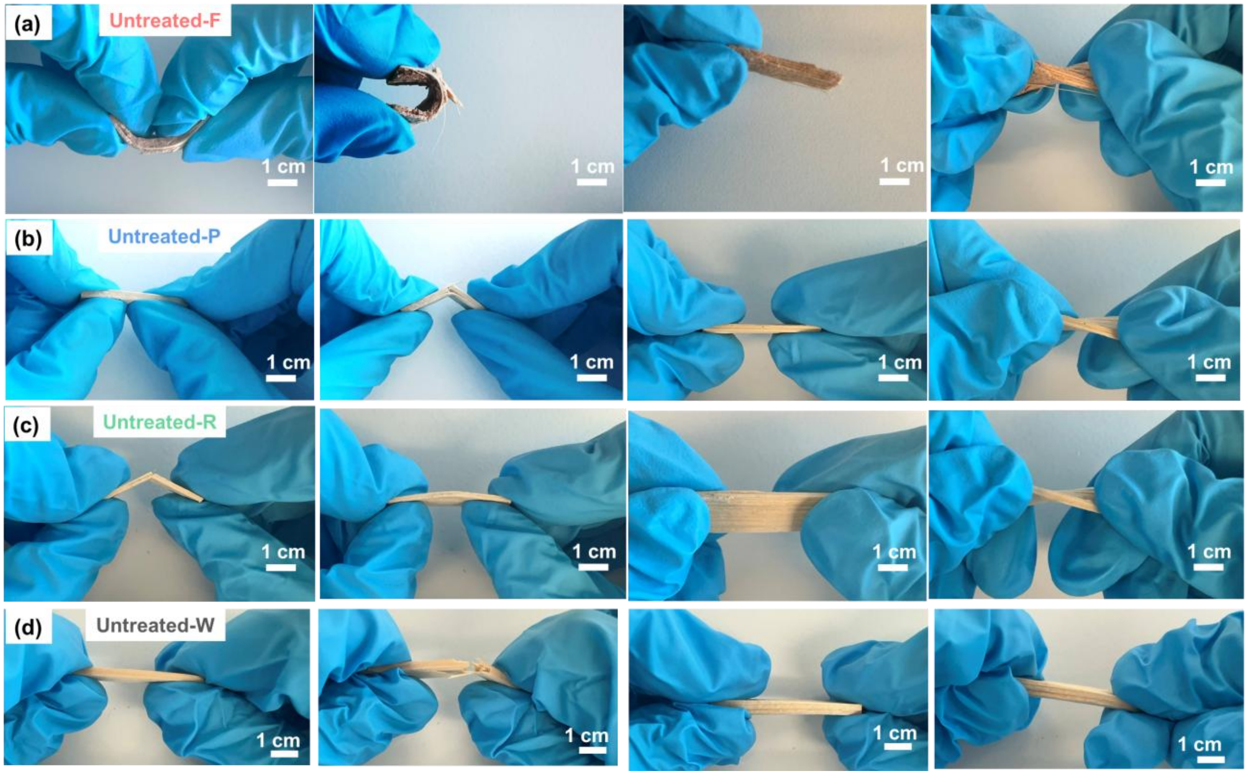

Figure S 4. Mechanical deformation behavior of untreated samples under manual bending of (a) Untreated-F, (b) Untreated-P, (c) Untreated-R, and (d) Untreated-W.

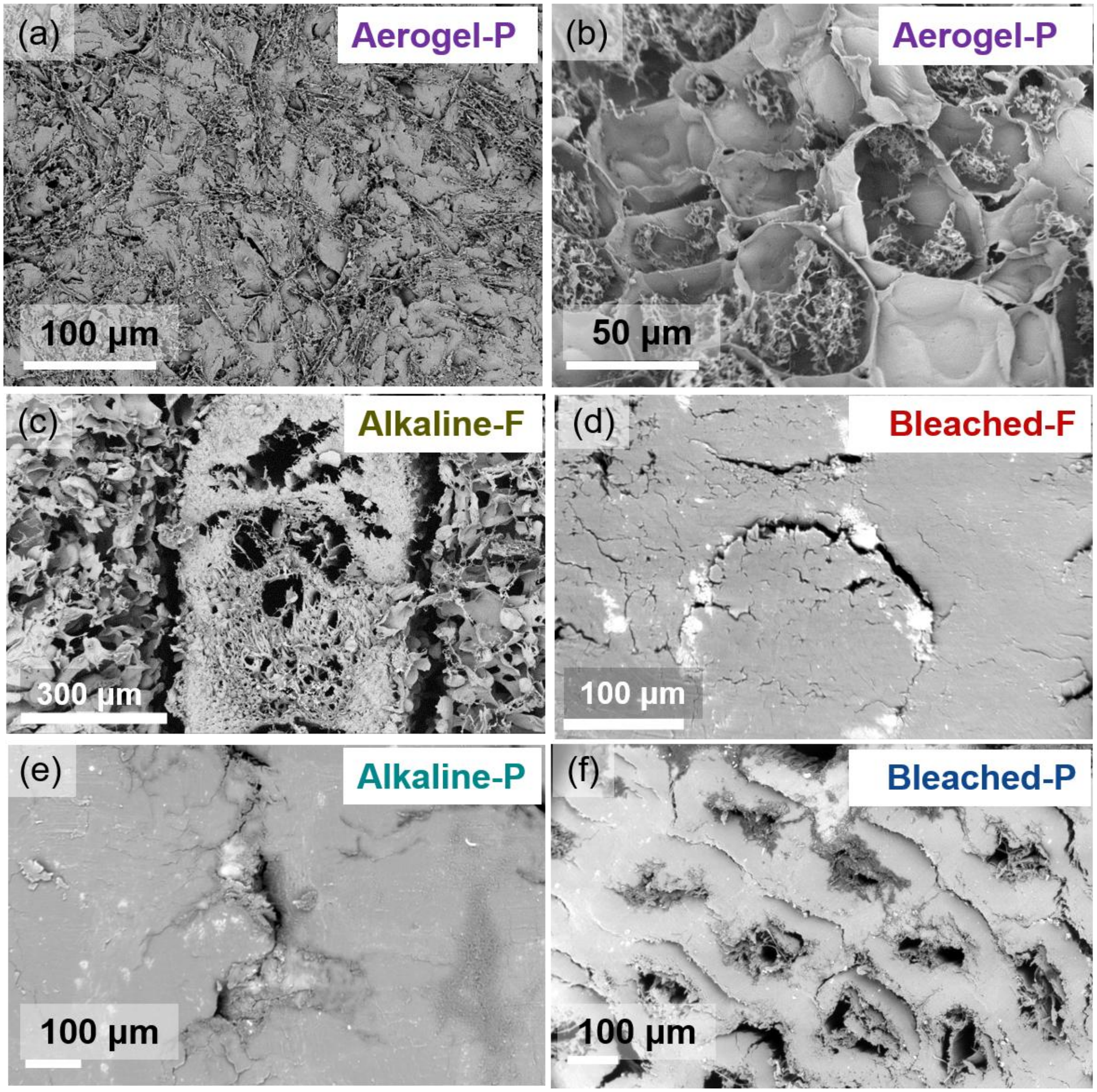

Figure S 5. SEM images of (a, b) Aerogel-P, (c) Alkaline-F after freeze-dring, and (d) Bleached-F, (e) Alkaline-P, and (f) Bleached-P after air-drying.

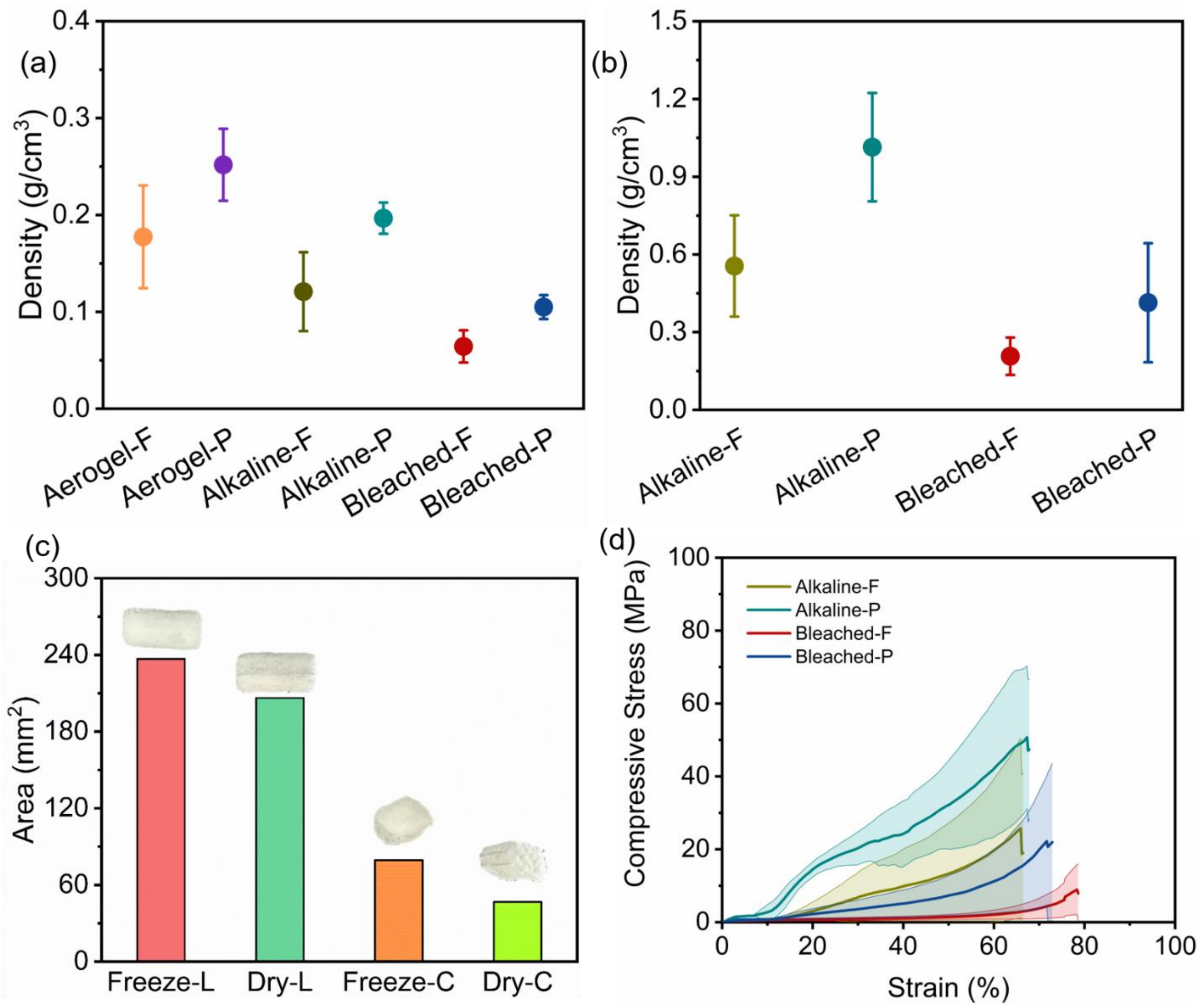

Figure S 6. Density of delignified samples by (a) freeze-drying, and (b) air-drying. (c) Aerogel-F area before and after freeze-drying in longitudinal direction (Freeze-L, and Dry-L) and cross section (Freeze-C, and Dry-C). (d) Mean transversal compressive strength curve of treated samples after air-dried.

### 2. Supplementary discussions

#### Supplementary Discussion 1-Chemical properties of untreated samples

The chemical compositions of untreated samples were evaluated by FTIR spectroscopy as shown in Figure S1 c, d. The stronger band near 3340 $cm^{-1}$ indicated a greater abundance of hydroxyl groups [1], while variations near 1730 and 1510 $cm^{-1}$ were associated with differences in hemicellulose- and lignin-related structures, respectively [2], [3]. Date palm-derived samples displayed a lower absorption band at approximately 1510 $cm^{-1}$ than Untreated-W (Figure S1d), indicating a lower abundance of aromatic structures in date palm-derived samples.

Thermogravimetric analysis (TGA) was performed in air to further elucidate the composition-dependent multi-stage thermal degradation behavior of untreated date palm and wood samples (Figure S1e). An initial minor weight loss below ~150 °C corresponded to the evaporation of physically adsorbed moisture. This was followed by a major mass loss between ~200 and 380 °C, associated with the decomposition of hemicellulose and cellulose. Compared to wood (Untreated-W), the date palm-derived samples (Untreated-F, -P, and -R) exhibited a slightly earlier onset of degradation because of their relative enrichment of hemicellulose content (Figure S1d). In the higher temperature range (~380-550 °C), a broader and more gradual mass loss was observed, corresponding to lignin degradation [4]. Notably, palm-derived samples displayed a more extended degradation profile, reflecting their structural heterogeneity. At temperatures above ~550 °C, the residual mass of palm materials remained higher than that of wood, suggesting a greater inorganic (ash) content typical of non-wood biomass [5]. Overall, these differences in thermal behavior highlighted the distinct compositional and structural characteristics of date palm tissues compared to conventional wood.
